# Valuing Carbon Stocks across a Tropical Lagoon after Accounting for Black and Inorganic Carbon: Bulk Density Proxies for monitoring

**DOI:** 10.1101/824490

**Authors:** John Barry Gallagher, Swee-Theng Chew, John Madin, Anitra Thorhaug

**Affiliations:** Institute for Marine and Antarctic Studies (IMAS), University of Tasmania, Hobart 7000; Borneo Marine Research Institute (BMRI), Universiti Malaysia Sabah, Jalan UMS, 88400 Kota Kinabalu, Sabah, Malaysia.; School of Forestry & Environmental Sciences, Yale University, New Haven, CT

**Keywords:** Blue carbon, Mangrove, seagrass, Salut–Mengkabong, Sama-Bajau

## Abstract

Managing seagrass and mangrove can be enhanced through carbon valued payment incentives schemes. Success will depend on the accuracy and extent of the carbon stock mitigation and accessible methods of monitoring and marking changes. In a relatively closed socioecological Southeast Asian lagoon we estimated the value of total organic carbon stocks (TOC) of both seagrass and mangroves. Mitigation corrections were also made for black carbon (BC) and calcareous inorganic carbon equivalents (PIC_equiv_), and their sediment dry bulk density (DBD) tested as a cost effective means of both estimating those stock concepts and possible impacts outside their parameter confidence intervals. Overall, seagrass and mangroves TOC densities across the lower lagoon ranged from 15.3±4.3 and 124.3±21.1 Mg C ha^-1^ respectively, 175.2±46.9 and 103.2±19.0 Mg C ha^-1^ for seagrass and 355.0±24.8 and 350.3±35.2 Mg C ha^-1^) for mangroves across the two upper lagoon branches. Only mangrove biomass made significant additional contributions ranging from 178.5±62.3 to 120.7±94.8 Mg C ha^-1^ for lower and upper regions respectively. The difference between the lagoons total seagrass and mangroves TOC stocks (5.98±0.69 and 390±33.22 GgC respectively) was further amplified by the lagoons’ larger mangrove area. When corrected for BC and PIC_equiv_, the carbon stock mitigation was only reduced by a moderate 14.2%. Across the lagoon the sedimentary DBD showed strong (*R^2^* = 0.85, *P* < 0.001) to moderate (*R^2^* = 0.67, *P* < 0.001) linear correlations with seagrass and mangrove [TOC] respectively, moderate correlations with seagrass [PIC] (*R^2^* = 0.6, *P* < 0.001), but an invariant and relatively constant response to mangrove [PIC] (2.7 kg m^-3^ ± 0.07). Valuations as CO_2_*e* was worth on average 0.44 million US$ y^-1^ over 20 years; less than the total income of the indigenous users as potential custodians (1.8 and 7.4 million US$ y^-1^). Implications of this valuation was discussed.

## INTRODUCTION

Coastal vegetated or blue carbon ecosystems of mangroves, salt marshes, seagrasses and seaweeds, support a range of ecosystem services that benefit both local users, and collectively can mitigate global greenhouse gas emissions (Nellemann, Corcoran *et al*. 2009). Measuring the suit of these natural capital services does not easily fit within traditional economic models of supply and demand of tangible goods and services (Costanza, de Groot *et al*. 2014). The advent greenhouse gas emissions, however, has led to the emergence of blue carbon mitigation investment schemes. These schemes do fit comfortably within a frame work of supply and demand of tangible goods (i.e., carbon stocks) and services (i.e., mitigation of greenhouse gases) (Hejnowicz, Kennedy *et al*. 2015; Pendleton, Donato *et al*. 2012). The tenet behind such schemes is to value the cost of damage to the global environment from the loss and disturbance of ecosystem carbon sinks. This is achieved through a generic form of carbon cap and trade on the open market. The cap on user carbon emissions is set by the policy within the economic region. Users that emit more than their cap can do so by purchasing carbon credits to the custodians and managers of those ecosystems as a payment scheme for this ecosystem service (PES). The value of credits is then ultimately determined by capacity of the ecosystem persistence to mitigate any further greenhouse emissions, at a price set by the market competition (Repetto 2013). The schemes thus provide both incentives for users to monitor and protect these ecosystems as well as for traders to reduce their payments and emissions below the cap. No more is this important than for coastal communities within SE Asia. Such communities have a stake in maintaining the integrity of seagrass and mangroves as their major supply of food, building materials, cooking fuel and disposable income (Raduan, Ariff *et al*. 2010). Clearly, PES requires good estimates for carbon biomass and sedimentary contents that statistically representative sampling can provide.

### Extent of the carbon stocks

While measurements of carbon within biomass and sediments can be relatively straightforward, questions have been raised on how stock preservation is assessed as a mitigating service in further release of the dominant greenhouse CO_2_ from the marine environment. That is, questions of the extent of loss after disturbance, how much is remineralised over climatic scales, whether to include the presence sedimentary allochthonous organic recalcitrants and particulate inorganic carbon (PIC) as an arbiter of carbon stock mitigation services (Chew and Gallagher 2018; Gallagher 2017). For woody biomass, in particular, how much is released as CO_2_ could vary with its use, should it be disturbed or harvested (Eong, 1993). For sediments the depth disturbance and fate of the sedimentary carbon stock will depend on the nature of the disturbance as a replacement ecosystem or land use (Siikamäki, Sanchirico *et al*. 2013)

The potential loss of seagrass biomass stock to remineralisation can be arguably linked to similar fate as its advected litter deposits, when we consider its utility as commercial product as limited. How much is reburied (Sophia and Robie 2016) and sequestered to the deep ocean is uncertain (Duarte and Krause-Jensen 2017; Gallagher 2015). This is in contrast the fate of mangrove biomass, where uncertainty is housed in assessing to estimate the fraction mineralised in the production and use charcoal over the amount stabilized as building materials and artesian products (Eong 1993).

For the sedimentary total organic carbon (TOC) stocks assessments must also be made of how much will be remineralised within climatic time scales. That will depend largely, not necessarily on the total depth of the sediment column, but the depth of disturbance and how much of the TOC over that depth is remineralised within climatic time scales (Pendleton, Donato *et al*. 2012; Siikamäki, Sanchirico *et al*. 2013). While, the condition ‘likely’ can be hedged (Gallagher 2017), it is ultimately unknowable. Consequently, IPCC (2014) have suggested allocation of precautionary limits on parameters of disturbance and remineralisation to move PES schemes forward. A likely depth of soil disturbance has been set to a maximum of 1 m, with remineralisation rate set at 96%. This scalar quantity is then given a vector characterization by averaging the loss to take 20 years to reach equilibrium (IPCC 2014). Although its applicability to coastal marine systems has not been fully considered and maybe not appropriate (Järviö, Henriksson *et al*. 2018; Pendleton, Donato *et al*. 2012; Thorhaug, Poulos *et al*. 2017). Nevertheless, the IPCC disturbance parameters are useful to temper carbon stock over-estimates that could lead to perverse outcomes in allowing traders to emit more than the system can mitigate.

### Exclusion of types of organic and inorganic carbon

Stable allochthonous organic forms, such pyrogenic black carbon do not require protection from mineralisation and thus, cannot be included in blue carbon ecosystem mitigation services (Chew and Gallagher 2018; Gallagher 2015; Gallagher, Chuan *et al*. 2019a). The exclusion or inclusion of particulate calcareous inorganic carbon (PIC) is not as clear. Disturbance of blue carbon sediments, the arbiter of how stock mitigation services are assessed, leads to carbonate oxidation and dissolution (Howard, Creed *et al*. 2017; Ware, Smith *et al*. 1992). This results in an increase in dissolved inorganic carbon (DIC) considered as a long term atmospheric sink (Maher, Call *et al*. 2018) and can theoretically reduce the water columns *p*CO_2_ below atmospheric pressure. This is provided the exchange neighbouring waters are sufficiently alkaline and /or its exchange restricted. In relatively warm tropical/subtropical waters, however, it has also been observed that sediments vulnerable to disturbance may also lead to whiting events. These events are the expression of water column calcareous carbonate precipitation. The mechanism is under some debate. Events are either seeding from sedimentary particles in waters already saturated with calcium carbonate salts, and/or forced by the additional increases in pH and alkalinity (Broecker, Sanyal *et al*. 2000; Sondi and Juračić 2010). The first mechanism suggests an increase in the molar fraction (0.63) of *p*CO_2_ equivalent for the carbonate precipitated (PIC_equiv_). The coefficient of 0.63 is a chemical speciation distribution determined by the water body’s alkalinity, pH, salinity, and the current atmospheric partial pressure (Ware, Smith *et al*. 1992). The second mechanism suggests that the additional carbon stock service provided by DIC from carbonate dissolution is constrained by re-precipitation. Until the science is convergent, we suggest that carbon stock assessments subtract PIC_equiv_, along with BC, from the total carbon stock. In this way, perverse environmental outcomes may be avoided when traders have effectively been given permission to emit beyond the capacity of the ecosystems’ carbon sink (Sophia and Robie 2016).

Once the carbon stock has been valued for trading, there is an obligation by the custodians to monitor this capital investment. While measuring biomass using its carbon parameters are well understood, the carbon stock variability within sediments would benefit from a predictor accessible to the local citizenry. Relatively Sophisticated element analysis of content, and cheaper more accessible laboratory organic matter and carbonate content combustion methods (Santisteban, Mediavilla *et al*. 2004) are beyond the capacity of fisher communities. The possible exception, are motivated school programs (Martay and Pearce-Higgins 2018) that have access to basic laboratory equipment supported by trained teachers or experts in the field. A simple cost effective and accessible alternative is of course preferable. In this way, it can conceivably be used to monitor the region fot the effects of local disturbance or upstream catchment developments, and continue to test the valuation by extending the assessment for areas not yet fully sampled. Dry bulk density (DBD g dry mass cm^3^) is one potential cost-effective measurement proxy for soil organic carbon. To date this has been limited to tropical peatlands (Warren, Kauffman *et al*. 2012), The authors found that DBD had an positive proportional response with peat soil organic carbon concentrations, but became increasingly unpredictably at high bulk densities equivalent to < 40% organic carbon concentrations.

As far as we are aware DBD relationship with carbon concentrations has not been formally expressed across tropical seagrasses and mangroves or indeed empirical relation to PIC concentrations. Nor is it likely that DBD would respond in the same proportional manner to organic carbon concentrations, as unlike peatland sediments, the DBD is dominated by its organic components. Leaving aside the degree of water saturation in intertidal conditions, for seagrass and mangrove sediments the DBD is largely increases with and increasing sand content over the smaller silt and clay components and their characteristic organic matter contents and larger interstitial pore water volumes (Tolhurst, Underwood *et al*. 2005). The mineral organic content is mostly associated with the smaller clay-silt fraction, the result of larger surface sorption area, and interestingly is invariant across differing soils and regions of the globe (Mayer 1994). Other contributions can come from the remaining mass and water content of the plant litter, along with any bioturbated channels or gas vacuoles (Tolhurst, Underwood *et al*. 2005). Consequently, unlike peat soils we should expect an inverse proportionality of DBD with organic carbon concentrations. The variance across the residuals then becomes a likely expression of differing net litter rention and possibly variance associated with surface bioturbation and vacuoles. Should the relationship of be useful for TOC concentrations, its established relationship with BC across the lagoon can be used to adjust the organic carbon fraction involved in mitigation services (Chew and Gallagher 2018; Gallagher, Chuan *et al*. 2019a).

For PIC, as far as we are aware however, no relationship with DBD has yet been tested or any suggested reason other than a useful and possibly lagoon specific empirical coincidence. Nevertheless, for the sake of rigor, it is postulated that PIC concentrations associated with epibiont production may be proportional to the density and size of the seagrasses’ canopy (Perry and Beavington-Penney 2005). Furthermore, this is covariant with increases in species canopy size up the lagoon, and a falling DBD with increasing proportions of the slit-clay fraction. In other words, PIC concentrations across this lagoon could be expected to increase with falling DBD in some manner. For mangroves the dynamic with PIC concentrations are mainly associated with epibenthic production and in proportion to the supply of food as mangrove litter (Volker and Matthias 2002). At worst, it is postulated that the remaining components that control the changes in DBD, and litter supply from changes in canopy biomass across the lagoon would likely confound any significantly relationship with PIC concentrations. At best, falls in biomass density with increasingly sandier and higher DBDs’ across the lagoon are sufficiently large to reduce the influence of other confounding factors.

### Aims

This study seeks to identify the variability, the total extent and PES annual monetary values of seagrass and mangrove TOC stocks from biomass and sediments across upper and lower regions of Salut–Mengkabong lagoon (Sabah, Malaysia). The values are then tested for future conceptual bias considerations by sequentially accounting for sedimentary BC and PIC*_equiv_*. Given BC can be methodological concept two divergent means of analysis were used to measure concentrations. Estimates of the PES values were then put into context. The total annual income generated by the indigenous fishers, current occupants of lagoon (Sama-Bajau), was adjusted to current rates, along the efficacy of the response of sedimentary TOC and PIC concentrations with DBD. In this way, the total value of the lagoons carbon stocks become a measure of potential for managerial incentives for custodians, and how it may be best applied.

## METHODS

### Study Site

This is a description of the Salut-Mengkabong lagoon, its general topography, climate, the type of coastal vegetated ecosystems, and population rural/urban character. Sampling techniques and storage are described along with determinant analysis, data processing, the sampling design and statistical methods used to assess carbon stocks concepts.

Salut-Mengkabong lagoon (6.101734°N, 116.153845°E), is a tropical tide-dominated system semi-rural region (Hogue *et al.,* 2010) situated several kilometers south of the district of Tuaran (population ∼100,000). The lagoon has two major branches, Salut and Mengkabong, served by a narrow but relative deep entrance (up to 6 m). Mangroves occupy its shoreline and its water body supports subtidal seagrass meadows along an ostensibly lower sandy and upper muddy sedimentary gradient (Figure 1). Development around the shoreline is moderate with some aquaculture ponds at the head of the Salut branch and inland of the north shore of the Mengkabong branch. Discussions with the appointed community head (2017) indicated that the lagoon supports 480 households of the Sama-Bajau community (unpublished data). Of the 480 households 96% of the community families were fishers that depend on the lagoons’ mangrove and seagrass resources (Raduan *et al*., 2010). While inputs of BC across the Southeast Asian region as a whole is substantial, it should be noted that Sabah lies only within the Penumbra of atmospheric smoke haze emanating from the distant Chinese mainland and seasonal Kalimantan peat fires (Permadi, Kim Oanh *et al*. 2018).

**Figure 1.**
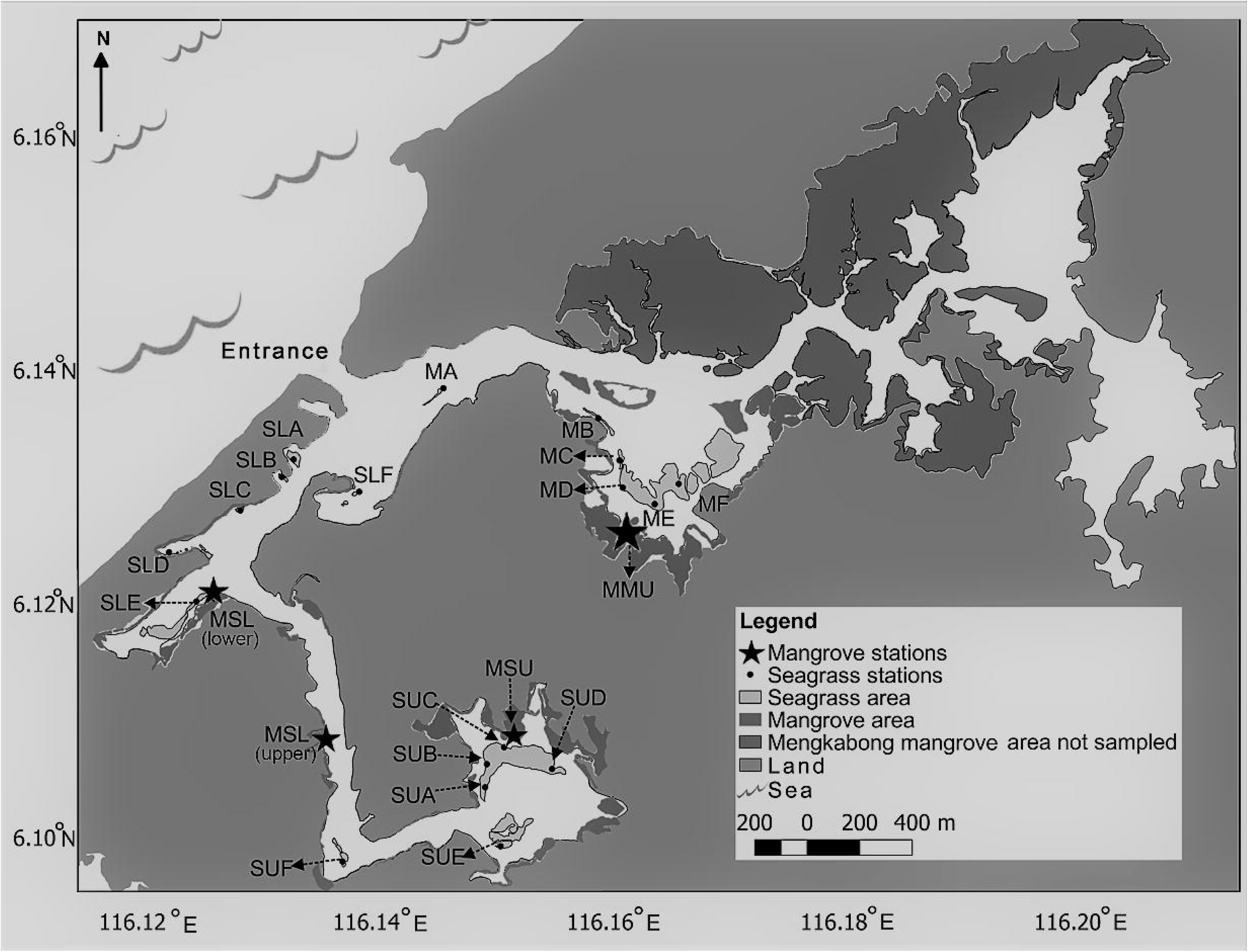
Mangrove and seagrass sampling stations. The stations are centered between replicated sampling sites and compiled across lagoonal regions. Seagrass stations within the lower and upper Salut branch are labelled, SLA, SLB, SLS, SLD, SLE and SUA, SUB, SUC, SUD, SUF respectively, and seagrass stations within the Mengkabong branch are labelled MA, MB, MC, MD, ME, MF. Mangrove stations within the lower Salut branch are labelled MSL(lower) and MSL(upper), for the upper Salut branch MSU, and for the Mengkabong MMU.

### Collection and transport of samples

This is a description of the method and scale of collection, transport, storage of seagrass and mangrove sediments, and means by which biomass was measured and calculated.

For seagrass, sediment cores around 25 cm long were extracted with a 5 cm diameter PVC tube (*n* = 56) at each station across all of the lagoons seagrass meadows (Figure 1). On extraction, the cores were then immediately placed under ice in a vertical position before being transported to the laboratory and stored at −20°C after sampling for dry bulk density and sediment particle size. Coverage was averaged from two quadrats (50 x 50 cm) placed around the coring station and based on species or species mix estimated in accordance with Seagrass Watch flash cards (McKenzie and Yoshida, 2011). The aboveground biomass, ash-free dry weight, AFDW g/m^2^, from a previous study (Ismail, 1993), were converted to total dry mass t ha^-1^, with the assumption that AFDW is 80% of the total dry mass (Duarte and Chiscano, 1999). Their carbon contents were then calculated as an average of 28% C dry wt^-1^, taken from the leaves of mixed bed meadows from the adjacent Sepanggar bay (Table S1 Supplementary Materials, along with their stable isotope of C and N).

For mangroves, transect lines were laid out at randomly selected stations (∼ 500 m apart) across both the Salut and Mengkabong lagoon branches (Figure 1). Transect sampling stations were placed every 25 m (50 to 100 m long), which ran perpendicular to the shoreline. Sediment cores around 50 cm long were taken with 11 cm larger diameter PVC tubes (*n* = 20 x 2). The larger diameter made core penetration easier and eliminated sediment compaction. Immediately upon extraction, the cores transported to the laboratory under ice, then and stored at −20°C after sampling for dry bulk density, pore water salinity and particle size analysis. The depth approximates half the carbon stock to 1 m (Chmura, Anisfeld *et al*. 2003) and was corrected accordingly. Biomass within each plot (10 m x 10 m), was taken from mangrove tree diameters, measured at chest height, DBH (> 1.3 m). The DBH measurements and wood density (ρ) for the different species (Duke, Mackenzie *et al*. 2013; Josue and Imiyabir 2014; Kauffman and Donato 2012) are required to calculate biomass using a Southeast Asian allometric equation (Kauffman and Donato 2012) of which around half (parameter 0.5, (1) was considered as organic carbon (IPCC, 2014).

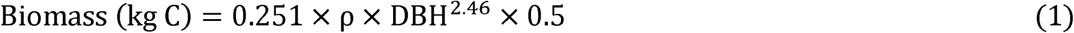

The area of the mangroves forests were incorporated with secondary data from Spalding *et al*. (2010), Google Earth satellite images 2017 and our ground truth information. For seagrass meadows, areas were measured by stepping around their subtidal outer perimeter at low tide, as guided by google earth images, and positions taken every few meters with a handheld GPS. In regions that were uncomfortably muddy and deeper, a boat was used to follow close to the meadows perimeter.

For seagrass around 25 cm of sediment were extracted with a 5 cm diameter PVC tube (*n* = 56) for sedimentary carbon stocks. The depth approximates half of the carbon stock to 1 m (Lavery *et al*., 2013) and placed immediately under ice in a vertical position before being transported to the laboratory. In each plot, two quadrats (50 x 50 cm) were used to study seagrass coverage according to Seagrass Watch protocol (Mckenzie and Yoshida, 2011). The coverage to biomass was obtained from previous studies on seagrass biomass in Salut-Mengkabong estuary/lagoon (Ismail, 1993).

### Sediment analsysis

All sediment dry bulk densities were measured from the homogenised core sample sealed within its plastic storage bag a sealed plastic bag. For seagrass, this was top 25 cm, and for mangroves the surface 25 cm and deeper core sections (25-30 cm and 25-40 cm) were processed separately. A cut off 1cm^3^ disposable syringe used in the manner of a piston core was used to measure the volume before the dry weight (Lavelle, Massoth *et al*. 1985), and after corrected for dissolved salt content using a refractometer from a subsample centrifuged to isolate its pore water. Subsamples were also taken from the homogenised sediments for sediment particle size distribution. A laser diffraction (LISST-Portable XR Sequoia Scientific) with full Mie theory in its calculations was used for the particle size spectrum, using a Wentworth classification from the clay/silt and sand proportions. For carbon analysis, the sediment was first shaken through 1 mm stainless steel mesh after drying (60 °C) and finally ground into fine powder (< 63 µm) with a porcelain mortar pestle. Subsamples were taken for PIC using a loss on ignition procedure (Santisteban, Mediavilla *et al*. 2004). Details of TOC and BC, as isolated by chemo-thermal oxidation (CTO) and after nitric acid oxidation can be found in Chew and Gallagher (2018). Again, all contents were reported after correction for salt content (Lavelle, Massoth and Crecelius, 1985).

### Carbon stock Analysis

The carbon stock densities and of top 25 cm and 50 cm for seagrass and mangrove respectively were calculated from fraction of the average TOC content, as its dry mass, down the length of the sediment core, its average bulk density and area of coverage (2).

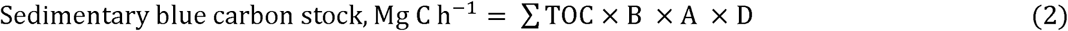

Where A is estimated area of coverage, B the dry bulk density, and D refers to the depth of the sediment core (Nellemann *et al*., 2009; Lavery *et al*., 2013; Howard *et al*., 2014). The carbon stocks for top 1 m were conservatively estimated by doubling up the storage from the 25 cm of the seagrass core (Fourqurean *et al*., 2012), and double the sum of the average top 25 cm and bottom 25 - 50 cm of the mangrove cores.

### Statistical analysis

A hierarchical random sampling design was used to estimate regional and lagoon wide mean and variance (standard error, S.E) of biomass and sedimentary stocks (Figure 1). For seagrass, scales were based on the determination of variability of independent seagrass sediment coring and biomass quadrat ‘stations’ from 2 to 4 ‘site’ replicates (10^1^ m scale) separated by the common length scale distance (>200 m) within a meadow and between meadows within lagoon regions. Regions were defined ostensibly by the sediment particle sizes, which trended from sandy within the lower regions to sandy-clay (Wentworth classification) within the upper parts of the lagoon (Table S2 Supplemental Materials): Lower upper Salut across the whole of the lagoon (10^3^ m). Upper Salut (SU) and Mengkabong (MU), and the lower common way region that extends to the entrance, named as Salut (SL) (Figure 1). With the exception of SL seagrass region, a Nested ANOVA structure was used to estimate the means of all variables and their summed variances across the estuarine regions. The number of degrees of freedom being reduced to the number of stations and number of sites within each region. This was calculated using the statistical functions and formulas within Excel™ on appropriate organized series column variables representing station and sites. For the SL region the means for TOC, BC and POC sedimentary variables, stations were weighted by area (0.5) between the largest southern meadow’s stations (SLE) and the remaining meadows. This was done to remove regional bias in the mean by a much larger sedimentary carbon stock density within the SLE meadow (10 fold greater) over the remaining meadows within the SL region (Figure S1 Supplemental Materials). For the sedimentary seagrass TOC-BC-PIC concept, direct calculations for comparisons to the previous true TOC-BC sampling population mean was confounded by an incomplete sets of stations for corresponding PIC stocks. In other words, while each stations’ TOC-BC variables necessary values falls because of subtraction of its corresponding PIC, it will not necessarily (or likely) change in a proportional manner from different estimates of sample means incomplete station sets. To normalize bias for direct comparisons with the TOC-BC sample mean, the percentage differences between means were calculated for the truncated TOC-BC-PIC stock with only its corresponding TOC-BC variables. This difference was then used to estimate a corresponding fall in the sample mean from the compete TOC-BC set to a now modelled mean of the truncated TOC-BC-PIC set. Variances for summation were then taken from the truncated set. The set was incorporated within the Nested ANOVA as variability about the new mean for the TOC-BC-PIC concepts. It could be expected that the variance of the truncated set was ostensibly equal or greater than a complete set, and thus remove any tendency towards a type one error when comparing sample mean between regions.

We refer the reader to Supplementary information and files for data that relate to the Figures, as well as additional tables and figures for parameters and smaller scale illustrations of stock density variance used for the weighting decisions for the Nested ANOVA.

## RESULTS

The section is description of the parameters that characterize the stations within the delineated regions of the lagoon, namely the plant species, seagrass coverage, the sedimentology. The information is used to constrain any stand out similarities or difference in carbon stock density concepts within and between regions. Finally, the total organic carbon stocks of seagrass and mangrove are calculated across the lagoon. The totals are valued in relation to estimates of the indigenous community’s annual incomes.

### Biological and physical parameters of seagrass and mangroves

All seagrass meadows across the lagoon were subtidal, with the visibly more turbid upper Salut and Mengkabong branches supporting the largest and monospecific *Enhalus acoroides* meadows (Figure 1). Both of these *Enhalus* sp. meadows were located near equidistant from the lagoons entrance but with discernibly greater canopy coverage for the Mengkabong meadow (Tables S3a,b Supplemental Materials). Five of 6 stations reported ≥ 50% coverage within Mengkabong with the majority of Salut meadows’ stations (i.e., site replicates) reported coverages of ≤ 50%. Across the remaining lower lagoon meadows, coverage was > 50% but with a mix of *Cymodocea* spp. and *Enhalus acoroides* species with an interchanging dominance. Of the mangrove canopy, *Rhizophora apiculata* was found to be the predominated tree species across the whole of Salut-Mengkabong lagoon with monospecific examples within the lagoons’ upper regions. This was in contrast to lower and middle sections of the Salut branches. Here, the stands were more diverse (*Rhizophora mucronata*, *Ceriops decandra* and *Lumnitzera racemose*) larger and seemingly more mature.

Overall, the subtidal sediments across the lagoon reflected the degree of isolation from the main channel and the distance from the lagoons’ marine tidal delta. The lower lagoon supported sandy sediments, but with a significantly greater clay-silt contents within the meadows at stations SLE and SLF (Figure 1) of 8.6 to 9.5% *cf* 0.9-2.5% (*P* < 0.001). For the upper Salut and Mengkabong regions, sediments ranged from sandy through to sandy clay loam as a function of distance from the marine tidal delta (Table S2 Supplemental Materials).

### Seagrass and mangrove carbon stock densities

The contribution of seagrass biomass to the TOC stock density across stations was restricted to upper region of the largest seagrass canopy and found to be insignificant and well within estimated sediment TOC stock density error (Table S3a Supplemental Materials). Consequently, for clarity, seagrass biomass was not explicitly included within the assessment analysis but expressed as less than other stock contributions’ lower significant figure

The lower Salut seagrass stock density concepts were found to be on average several times smaller than the remaining upper regions of the Salut and Mengkabong branches (*P* < 0.001) (Figure 2a, 2b and 2c). However, the relative difference within each region between concepts was not consistent. There was a different pattern of ranking and relative differences between stock concepts (Figure 2a, 2b and 2c). The upper Salut region, serviced by a small river (Figure 1), revealed a clear and greater successive fall in the concepts’ sample means over the lower Salut and upper Mengkabong regions. The differences originating from the larger BC stocks (Figure 2a, bars 2 and 4), in particular BC as isolated by CTO (Figure 2a, bar 4), and PIC stocks (Figure 2a, bars 3 and 5). This appeared to be the result of a disproportionate amount of BC, as separated by CTO (Figure 2d, bar 4) that could not spate the inclusion of photoliths within the concept (Chew and Gallagher, 2018).

**Figure 2.**
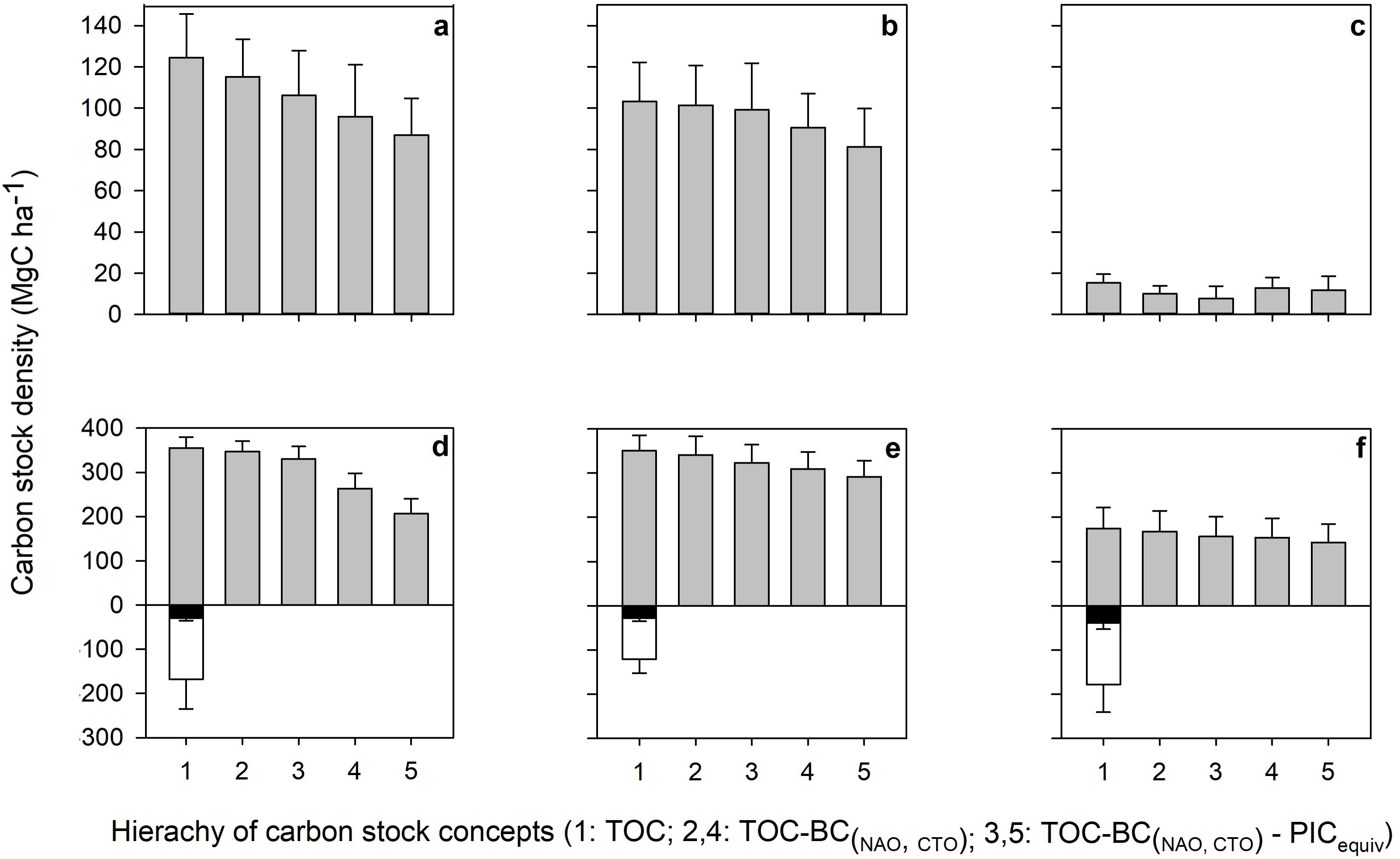
Sediment and biomass carbon stock density concepts for seagrass (a,b,c) and mangroves (d,e,f). Stocks are compiled within regions of the upper Salut (**a** and **d)** and Mengkabong branches (**b** and **e**), and the lower Salut branch (**c** and **f**). The grey bars represent each regions average sediment stock concepts calculated to 1 m depth: TOC (bar 1), TOC-BC_(NAO)_ (bar 2), TOC-BC_(CTO)_ (bar 3), TOC-BC_(NAO)_-PIC_(equiv)_ (bar 4), TOC-BC_(CTO)_-PIC_(equiv)_ (bar 5). The white bars and the contained black bars (d, e, f) represent the above below ground mangrove biomass respectively. The variance about their means are the standard errors taken from a Nested ANOVA of sites and station replicates.

In contrast, to seagrass, the variability of mangrove sedimentary carbon stock density concepts across regions were notably smaller, with means more than twice that of seagrass (Figure 2d, 2e and 2f) and irrespective of the differences in mangrove species (Tables S3a,b Supplemental Materials). The possibly exception was a close equivalence between TOC-BC_CTO_-PIC_equiv_ concept between the upper and lower Salut region (Figure 2d, 2f bars 5; *P_(same)_* = 0.015). Unlike seagrass, the mangrove biomass held a measurably significant fraction (Figure 2d, 2e and 2f) on average 47.4%, 34.5%, and 101.25 of their sedimentary TOC stocks for the upper Salut, Mengkabong and lower Salut regions respectively. The pattern also reflecting the below ground biomass contributions (Figure 2d, 2e and 2f), ranging around 17.5% to 22.2% of their total biomass. Like seagrass, all carbon stock density concepts were notably greater (*P* < 0.05) in the upper branches (Figure 2d and 2e) of the lagoon, although less so, than found in the lower Salut region (Figure 2f). Again, the larger fall in stock density concept services was found in the upper Salut region, the result of an apparently larger BC components, particularly as isolated by CTO (Figure 2e, bar 4).

### Total seagrass and mangrove carbon stocks for the lagoon

The values of the traditional TOC stock concept show that most of the carbon stocks are located within mangroves (Table 1). This is in part because of the mangroves’ greater carbon stock densities (Figure 2), but mainly because of the extensive, and relatively inaccessibly mangrove forest along and back from Mengkabongs’ north shore (Figure 1). On average 321.1 GgC appeared to be stored within this region (calculated as the aerial fraction of its Mengkabong TOC stock; (681.92/750.12) x 353.3, Table 1). The lagoons’ remaining TOC stocks, on average 8.9% (calculated from Table 1), is located within more accessible seagrass and coastal fringe mangrove forests. Overall, this variance is reflected in the other stock concepts as seen from their similar relative stock densities (Figure 2). If we focus on the stock concept with the greatest impact on traditional TOC stock estimates (i.e., TOC-BC_CTO_-PIC_equiv_), the degree of bias in the mitigation of greenhouse gas emissions 14.2% ± 0.12 (calculated from Table 1), appear as only a moderately smaller than traditional assessments.

**Table 1.** Total organic biomass and sedimentary carbon stocks within regions of the Salut–Mengkabong lagoon (Sabah, Malaysia). Corrections have also been made for exclusions of largest recalcitrant black carbon fraction (BC) as isolated by thermal oxidation (CTO), and the inorganic carbon concept PIC_equiv_. Variances represent nested standard errors compiled across regional stations and sites. Sgr and Mgr refer to seagrass and mangroves respectively.

| Region and ecosystem | Area (ha) | TOC sediment stocks (GgC) | TOC Biomass stocks (GgC) | Total TOC stocks (GgC) |
| --- | --- | --- | --- | --- |
| Upper Salut Sgr | 23.05 | $2.87 \pm 0.49$ | $< 0.01$ | $> 2.87 \pm 0.49$ |
| Mengkabong Sgr | 29.04 | $3.00 \pm 0.55$ | $< 0.01$ | $> 3.00 \pm 0.55$ |
| Lower Salut Sgr | 7.39 | $0.11 \pm 0.03$ | $< 0.01$ | $> 0.11 \pm 0.03$ |
| Upper Salut Mgr | 48.43 | $17.2 \pm 1.2$ | $8.15 \pm 3.3$ | $25.35 \pm 3.51$ |
| Mengkabong Mgr | 750.12* | $262.76 \pm 24.1$ | $90.54 \pm 22.5$ | $353.30 \pm 32.97$ |
| Lower Salut Mgr | 32.52 | $5.70 \pm 0.15$ | $5.80 \pm 2.03$ | $11.50 \pm 2.04$ |
| Total for Sgr | 59.48 | $5.98 \pm 0.69$ | $< 0.01$ | $> 5.98 \pm 0.69$ |
| Total for Mgr | 831.07 | $285.66 \pm 24.13$ | $104 \pm 22.83$ | $390 \pm 33.22$ |
| $\Sigma$ Total Mgr & Sgr | 890.55 | $294.63 \pm 24.17$ | $104.50 \pm 22.82$ | $399.14 \pm 33.20$ |
| Total TOC-BC<sub>CTO</sub>-<br>$PIC_{equiv}$ (Sgr & Mgr) | 890.55 | $237.86 \pm 25.15$ | $104.50 \pm 22.82$ | $342.36 \pm 33.96$ |

\*The area of the north shore Mengkabong mangroves (681.92 ha) is included in the stock calculations using the southern shore mangrove stock densities as representative of the region.

### Total organic carbon stocks values and the indigenous community income

Market values of carbon stocks are uncertain and depend on the choice of regional carbon credit market. Whether it is amenable or easily available, there is also uncertainty in any assessments on future price projections (Lavery *et al*., 2013; Siikamaki *et al*., 2013; Zarate-Barrera and Maldonado, 2015). For this study, we have chosen conservative estimates based on volunteer markets, currently 0.1– 0.02% of the value and volume of the regulated global carbon market. These are more flexible and known to fund micro-projects accessible to communities. Micro-projects are set at around US $10 per tonne of CO_2*equiv*_ (Peters-Stanley and Yin, 2013), but more commonly around US $6 per tonne of (Ullman, Bilbao-Bastida and Grimsditch, 2013). Converting carbon stocks to CO_2*equiv*_, results in total blue carbon stock credits traded at 1405.44 ± 116.93 GgC, and valued around 8.43 million US$. This one time scalar quantity can be transformed to average annual incomes as 0.44 million US$ y^-1^ over 20 years. The estimate assumes that, in general, the loss of stocks over this time period has likely reached equilibrium where 96 % of the carbon has been lost to the atmosphere (IPCC, 2014). When this estimate is corrected, for what we consider, as the most conservative carbon stock concept (TOC-BC_CTO_-PIC), both stocks and subsequent income would be only reduced by a moderate 14.2% across the whole lagoon. Nevertheless, for this type of lagoon, located within this Southeast Asian region, 14.2% is greater than traditional stock variability estimates (8.3% one tailed *P* < 0.05). Consequently, it can still be regarded as an important source of bias if not accounted for, and a bias that becomes increasingly important in more open coastal seagrass blue carbon ecosystems of the same region (Gallagher <u>et al., 2019)</u>

The household income of the lagoons’ Sama–Bajau community was estimated from an older estimated annual income of 12 000 to 48 000 RM per family (Raduan et al., 2010). The estimate was corrected for 2016 purchasing power as equivalent to 3 840 to 15 385 US$ y^-1^. The average exchange rates between 2010 and 2016 were used after correcting for the average rate of inflation experienced by Malaysia from 2011 to 2016. (https://www.poundsterlinglive.com/bank-of-england-spot/historical-spot-exchange-rates/usd/spots/USD-to-MYR; https://knoema.com/atlas/Malaysia/Inflation-rate). When scaled up to todays 480 households, taken from an interview with the Headman, the community is likely to have earn between 1.8 to 7.4 million US$ y^-1^.

### Bulk density as predictors of TOC and PIC sediment concentrations

Overall, the range of TOC concentrations within seagrass sediments were greater than found in mangroves, reflecting mainly the lower organic contents contributions from the sandier sediments (Figure 3). Over these ranges, there were strong to moderate correlations from least square regressions of DBD with concentrations for TOC, within the seagrass and mangrove sediments respectively, and for PIC within seagrass sediments. In contrast PIC concentration were both invariant with DBD and relatively constant (2.7 kg m^-3^ ± 0.07)

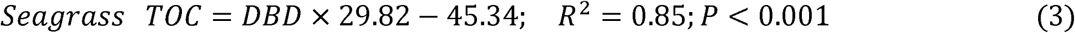

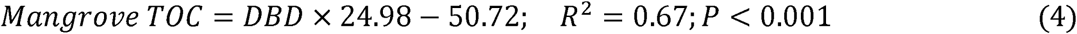

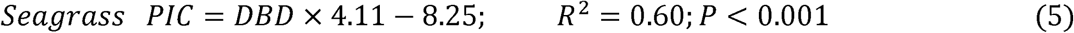

**Figure 3.**
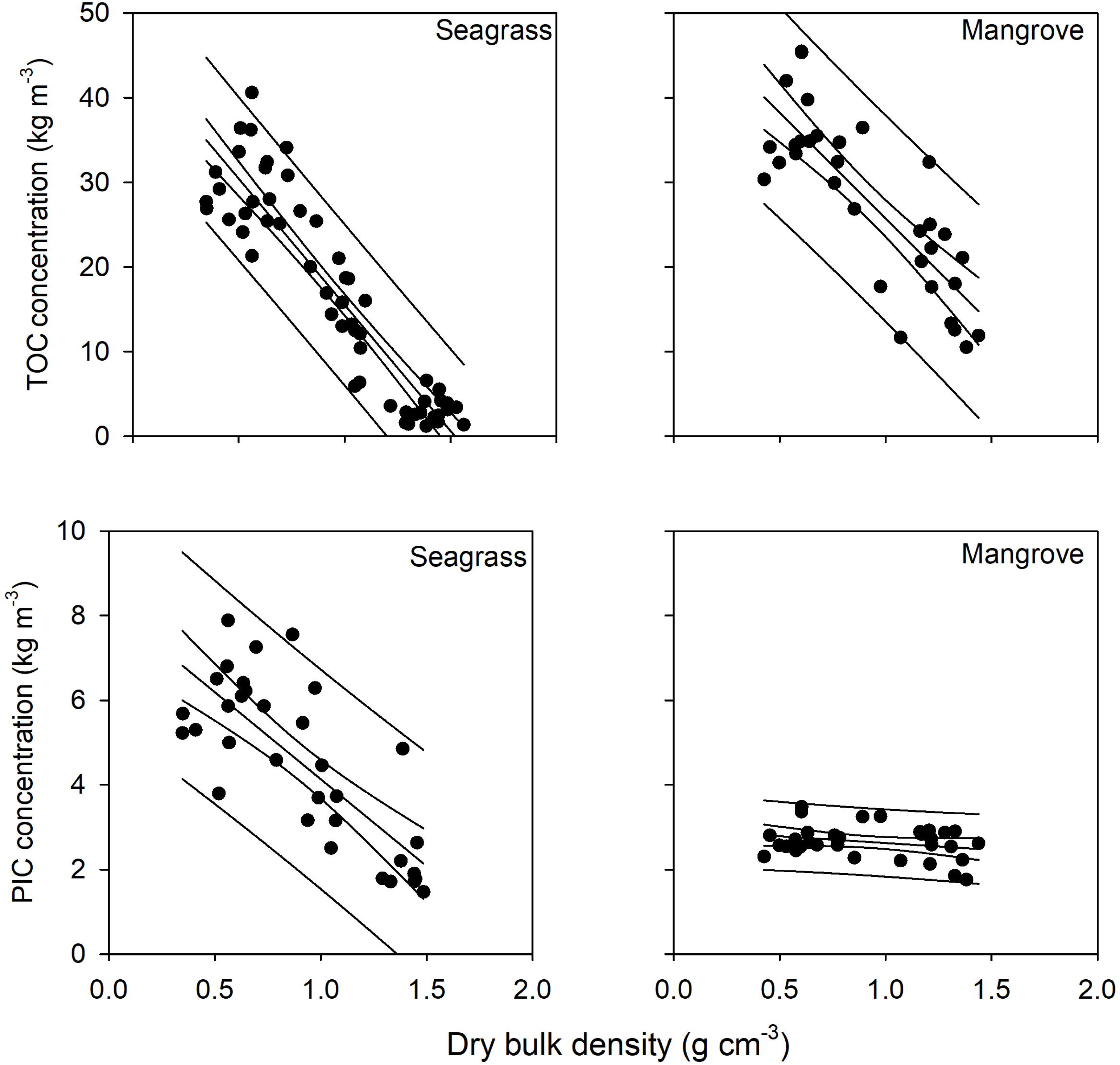
Dry bulk density relationships with sedimentary and [PIC]. The inner and outer statistical limits represent the 95% confidence limits and 95% prediction intervals respectively about their OLS regressions.

## DISCUSSION

Across the lagoon, it was clear that both seagrass and mangroves of the lower lagoon region supported smaller sedimentary carbon stocks. Explanations likely come from two standpoints. Firstly, greater net loss of litter and allochthonous particulates than the more sheltered upper lagoon embayments (Chiu, Huang and Lin, 2013; Portillo, 2014; Ricart, Perez and Romero, 2017; Gallagher *et al*., 2019a) The region is characterized by a typically more turbulent marine tidal delta and what it was observed as relatively fast flowing tidal flows along the narrow channel that lead towards upper part of the Salut branch (Figure 1). Secondly, it appears that in addition to allochthonous litter more than two thirds of sedimentary organic matter in seagrass sediments comes from the more extensive mangroves of the upper lagoon. It is recognised that other factors can also conceivably contribute to the variance. For seagrass, the size of the seagrass canopy, its coverage and the area of the meadow itself, are possible variants. The larger canopy species, as found in the upper lagoon, are more effective in retaining litter and contributing greater amounts to sediment deposition than the smaller faster growing species, which characterize the lower lagoon (Gallagher *et al*., 2019b). Larger meadows, such as found in the upper lagoon, appear to retain more litter by virtue of the main body having a greater geocentric distance from the edge (Ricart, Perez and Romero, 2017). Although, there is some suggestion of possible cofounding from the larger lower lagoon meadow at site SLE (Figure 1). Here the carbon stock density was notably larger than the remaining lagoons seagrass meadows (Table S3a Supplemental materials).

### Comparisons with other Southeast Asian traditional organic stock densities

Across the western quarter of the Southeast region, the lagoon’s seagrass total stock densities to 1 m of sediment are in general agreement to one another. For seagrasses of Chek Jawa, Singapore, within the shelter and embayment of the Johor straights, the carbon stocks were measured at around 138 ± se 8.6 Mg C ha^-1^ of (Phang, Chou and Friess, 2015). The island estuaries/lagoons of the Indonesian Archipelago to the immediate south of Borneo were measured at around the 129.9 ± se 9.6 Mg C ha^-1^, and 24 Mg C ha^-1^ within the more open turbulent coastal systems of its Pacific side (Alongi *et al*., 2016). It is only within the eastern sector of Southeast Asia, SE Sulawesi and SE Kalimantan (Borneo) do we find examples of notably larger carbon stock densities between 239.2 ± 44.9 (to 0.5 m) and 214.4 ± 48.7 MgC ha^-1^ (to 1 m) (Alongi *et al*., 2016).

Many previous studies of mangrove carbon stock densities have reported estimates to various depths (1 to 3 m). For comparison, we normalised the reports to 1m by a simple division. Similar to that for seagrasses, we found that the western section of Southeast Asia mangrove stock densities were similar to Salut–Mengakabongs’ upper lagoon. Chek Jawa’s mangrove stock density in Singapore was measured around 307 ± s.e. 33Mg C ha^-1^. The Indonesian Borneo (Kalimantan) mangrove forest recorded stock densities of around 356.51 ± s.e. 27.60 Mg C ha^-^1, with the Neighbouring Java mangrove stock densities were marginally lower (284.93 ± s.e. 15.62 Mg C ha^-1^) (Donato *et al*., 2011; Murdiyarso *et al*., 2015; Phang, Chou and Friess, 2015; Alongi *et al*., 2016). Like seagrass region stock density distribution noticeable greater stock densities were found in eastern sector of Southeast Asia; Sulawesi (759.07 ± s.e. 116.75 Mg C ha^-1^), Sumatra (542.81 ± s.e. 15.71 Mg C ha^-1^), and Papua (510.89 ± s.e. 81.06 Mg C ha^-1^) (Donato *et.al*., 2011; Murdiyarso *et al*., 2015; Alongi *et al*., 2016). Many of these mangrove ecosystems are located in more complex environs. Sulawesi mangroves grow nearby degrading coral reef, the result of eutrophication from high intensity farming activities (Alongi *et al*., 2016). Indeed, additional supplies of inorganic nitrogen may not only add to the biomass and sedimentary stocks through an increase productivity and leaf fall in these generally nitrogen limit systems, but add to sedimentary stocks through slow rates of soil organic mineralisation with increasing inorganic nitrogen supply (Fog, 1988). Other possible factors that distinguish this eastern sector are mangrove forests located next to a source of high carbon particulate runoff from freshwater peat swamp forests. However, unlike the mangroves of the region, the position of seagrass meadows are not ideally placed for trapping organic matter from freshwater peat swamps. Nor do seagrass productivity, in general, respond well to excessive eutrophication or turbidity (van der Heide *et al*., 2011). Nevertheless, moderate supplies of nutrients are capable of stimulating seagrass productivity particularly in acidic waters may stimulate seagrass productivity (Ravaglioli *et al*., 2017). In actual fact, these are the conditions seen within the eastern sectors of this archipelago where the Indonesian flowthrough nutrient upwelling dominates (Ayers *et al*., 2014). Here the origin of the deeper pacific equatorial current, rich in dissolved CO_2_ and nutrients, has a 30 cm pressure head over the Indian Ocean. This creates a major flow through in the eastern islands on the Sunda Plate (Wyrtki, 1987). In addition flows stream past Northeast Borneo from the China Sea. Together, continual supply of rich pacific equatorial water into the area in general, and with it substantial tidal exchange into estuaries and lagoons in it path. Indeed results throughout Southeast Asian region seem to reflect this (Thorhaug *et al*., unpublished, in review).

### Carbon stock carbon concept bias and variability

The seagrass meadows of the upper Salut branch supported significant contributions of sedimentary PIC, allochthonous BC and other possible recalcitrants. However, the dominance of mangroves carbon stocks across the lagoon and their associated smaller contributions from PIC across both ecosystems and BC in mangroves (Figure 2d, 2e and 2f) reduced the overall carbon stock services to a significant but moderate bias (around 14%). For BC factors controlling the size of its contribution across the lagoon has previous been tested by Chew and Gallagher (2018). The relatively small BC contributions to the total organic matter was in proportion to the total sedimentary organic carbon within seagrasses and relatively invariant in mangroves. The small contribution of BC as the result of relatively high rates of net productivity and deposition, and the BC delivery mechanism, dominated by soil wash out to seagrasses and the atmosphere to mangroves. For PIC, the small concentrations are in accord with the contention that for non-edaphic geological carboniferous sediments, dissolution may play a factor in moderating PIC stock densities (Saderne *et al*., 2018).

The relative variability of sedimentary PIC concentrations showed similar proportional and invariant responses across seagrass and mangrove stations as BC to organic matter, as inferred from its proportional relationship with DBD. It may be that there is an overlying variability in the supply of biological calcareous tests and shells to the surface sediments, associated with the changing seagrass canopy architecture, coverage (Perry and Beavington-Penney, 2005) and biomass. Across the lagoon, these factors are, in part, covariant with differing amounts of litter retention, their clay-silt fractions and organic contents that ultimately determines the changes in DBD. How PIC concentrations remain relatively constant in mangroves while independent of DBD determined by the differing supply of edaphic clay, silts and sands is not clear. Evidently, sedimentary PIC may require a simpler supply construct other than deposition of settling particles and subsequent burial. In place, we contend that PIC concentrations, as a minor fraction, become independent of DBDs’ components’ depositional supply when produced within a relatively constant surface bioturbation zone. In other words, the mass of detrital PIC saturated from the left overs of a burrowing benthic epifauna is a function of volume of the niche, and not its remaining mineral and organic composition. Not only within the current surface bioturbation zone but also a past memory of the bioturbation zones over depositional time.

### Dry bulk density: A tool for carbon stock concentrations

For both seagrass and mangrove [TOC] were respectively strongly and moderately inversely linearly related to DBD. Although, within the more muddy upper lagoon seagrass meadows the variance was greater. The greater amount of variance may reflect the variability in litter retention from the edge to the center of these larger meadows. As a management tool for additional measurements the correlation coefficient population confidence intervals (95%) appear to be very acceptable and well within 15% of the variability of many analytical methods (Byers, Mills and Stewart, 1978), let alone the lagoons sampling site variability. The latter variability is captured by their regressions’ prediction intervals. Indeed, this could be a useful boundary to monitor changes. For examples in changes in expected range of DBD densities carried out by the local community at for various stations. Also, as part of a more considered elemental organic carbon analysis targeted at those stations. That is to say, whether the response sits within the regression prediction limits as a first order assessment of change as either natural ecosystem variability or possible anthropogenic disturbance.

### Values of carbon stocks as a sufficient incentive for community management

It was clear that conservative estimates of the lagoons’ carbon stocks worth (0.44 million US$ y^-1^) was not sufficient to replace the income for every household of Sama–Bajau community (1.8 to 7.4 million US$ y^-1^). The significance of such a conservative valuation as an incentive for community management, however, may necessarily depend not on the size of population and their total income. The Sama-Bajau have an elected hierarchy that directs community sharing from allocated council grants for village community projects) as well as a sense of family income independence (Miller, 2011). The possibility of carbon trading in these circumstances then becomes not an alternative livelihood, but a way to augment income for themselves and/or benefit the whole community. Indeed the potential for both commitment and ability to conserve and monitor the lagoons’ coastal canopy and take sediment samples for future DBD measurements was evident from Mr Awang. Mr Awang was our teams’ boatman of the same Bajau-Sama community (see Acknowledgments), and has communicated concerns on the state and importance of the lagoons’ seagrass beds for his livelihood over a number of years to our team

## CONCLUSIONS

Traditional organic carbon stock densities for both seagrasses and mangroves across the lagoon are in broad agreement with other studies carried out in the eastern sector of Southeast Asia. The only notable exception are the smaller mangrove stock densities of the lower lagoon, also reflected in its seagrass stocks, and with other seagrass examples across the coastal eastern Southeast Asian sector. Accounting for more sophisticated carbon stock concepts by excluding BC and PIC_equiv_ contributions identified only moderate bias with traditional stock concepts, when integrated across the lagoons ecosystems (on average 14.2%). The use of DBD as a predictor of TOC and PIC concentrations or ecosystem parameter was found to be useful for both monitoring and potentially identifying future impacts. The conservative valuation of the lagoons’ total stock, as an annual income was not sufficient to totally replace indigenous community incomes. Nevertheless, we suggest that the potential exists for the estimates of 0.44 million $US y^-1^ to be directed towards improvements in the fisher community infrastructure, funds for training, or additional income for environmentally aware fishers within the community.

## AUTHORS’ CONTRIBUTIONS

JBG conceived the project, led the manuscript writing and the statistical analysis. STC designed the field sampling program, carried out measurements and collections, analysed the samples, and contributed to the manuscript and statistics. JM was in part responsible for the Supplementary Materials and contributed to project supervision. AT constructed reasons behind the high blue carbon stock densities unique to Southeast Asia and the regions’ information limitations. All authors read, copy–edited and approved the final manuscript.

## ACKNOWLEGEMENTS

We would like to thank the University Malaysia Sabah for providing funding under Grant SBK0239-STWN-2015. Thanks to all the volunteers includes Cheong Kai Ching, Yap Tzuen-Keat, Jason Lai Siang Kang, Lee Yin Lin, Erik Bin Naim and undergraduate students for helping out in sampling and laboratory work. Appreciation goes to our local Sama-Bajau boatman Mr Awang who shared his knowledge of Salut-Mengkabong lagoon and assisted in the sampling.

