## Supplementary Materials for "Valuing Carbon Stocks across a Tropical Lagoon after Accounting for Black and Inorganic Carbon: Bulk Density Proxies for monitoring"

**DESCRIPTION**

This document contains additional figures a cited in the main article for available biological and sedimentological parameters, and data used to present and calculate values of carbon stock concepts across sites and stations within regions of the Salut–Mengkabong lagoon (Sabah, Malaysia).

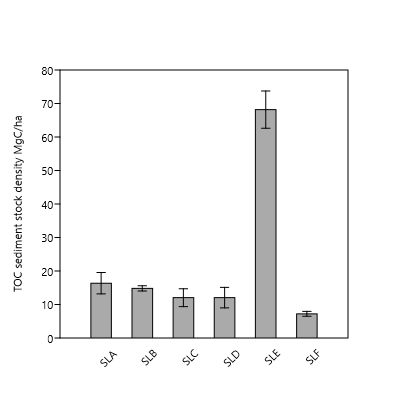

Figure S1. The unweighted mean and variability (s.e. n = 3) of sites within the lower Salut seagrass meadow stations of Salut–Mengkabong lagoon (Sabah, Malaysia). The figure is designed to illustrate the need for a interregional weighted average because of larger stock density outlier around SLE with the remaining stations in separate meadows (*P* < 0.0001). See Figure 1 in the main article for position of meadow stations.

Table S1. **Stable isotope and carbon and nitrogen contents leaf shoots collected around the vicinity of Gaya Island and Sepanggar Bay (Kota Kinabalu, Sabah , Malaysia )**. Analysis was performed after drying then fuming HCl at The Stable Isotopes in Nature Laboratory (SINLAB) Biology Department at the University of New Brunswick in Fredericton, New Brunswick, Canada.

| **Location** | **Date** | **Species** | **d13C** | **d15N** | **%C** | **%N** | **C/N** |
| --- | --- | --- | --- | --- | --- | --- | --- |
|  | 10/6/2016 | *Cymodocea rotundata* | -9.24 | 6.50 | 19.09 | 1.19 | 15.99 |
| Gaya Bay | 18/08/2016 | *Cymodocea rotundata* | -9.09 | 6.06 | 32.16 | 2.22 | 14.51 |
| Gaya Bay | 21/9/2016 | *Cymodocea rotundata* | -9.41 | 6.16 | 23.72 | 1.65 | 14.41 |
| Malohom Bay | 11/9/2016 | *Cymodocea rotundata* | -9.22 | 4.79 | 38.59 | 3.01 | 12.80 |
| Marine Ecology Research Centre | 22/7/2016 | *Cymodocea rotundata* | -10.09 | 4.91 | 33.70 | 1.84 | 18.34 |
| Marine Ecology Research Centre | 9/9/2016 | *Cymodocea rotundata* | -8.80 | 6.64 | 36.72 | 2.75 | 13.35 |
| Odec beach (UMS) | 16/6/2016 | *Cymodocea rotundata* | -9.73 | 5.48 | 21.47 | 1.34 | 16.04 |
| Gaya Bay | 10/6/2016 | *Cymodocea serrulata* | -11.40 | 7.09 | 26.41 | 1.62 | 16.32 |
| Gaya Bay | 18/08/2016 | *Cymodocea serrulata* | -11.03 | 7.05 | 23.34 | 1.45 | 16.12 |
| Gaya Bay | 21/9/2016 | *Cymodocea serrulata* | -12.47 | 6.98 | 37.90 | 2.60 | 14.57 |
| Gaya Bay | 11/12/2016 | *Cymodocea serrulata* | -11.79 | 6.40 | 26.87 | 1.61 | 16.70 |
| Malohom Bay | 22/7/2016 | *Cymodocea serrulata* | -10.27 | 4.67 | 22.77 | 1.36 | 16.77 |
| Malohom Bay | 11/9/2016 | *Cymodocea serrulata* | -10.54 | 5.62 | 37.89 | 2.29 | 16.52 |
| Marine Ecology Research Centre | 22/7/2016 | *Cymodocea serrulata* | -11.64 | 6.56 | 26.26 | 1.66 | 15.70 |
| Marine Ecology Research Centre | 9/9/2016 | *Cymodocea serrulata* | -9.90 | 6.68 | 37.24 | 2.12 | 17.54 |
| Malohom Bay | 22/7/2016 | *Enhalus acoroides* | -9.12 | 5.55 | 17.40 | 0.97 | 17.93 |
| Marine Ecology Research Centre | 22/7/2016 | *Enhalus acoroides* | -8.93 | 5.27 | 22.44 | 1.16 | 19.42 |
| Odec beach (UMS) | 16/6/2016 | *Enhalus acoroides* | -7.73 | 6.10 | 22.54 | 1.02 | 22.11 |
| Odec beach (UMS) | 16/6/2016 | *Halophila ovalis* | -8.62 | 5.98 | 27.01 | 1.88 | 14.41 |
| Odec beach (UMS) | 16/6/2016 | *Halodule uninervis* | -9.64 | 6.56 | 40.20 | 2.41 | 16.70 |
| Gaya Bay | 15/4/2016 | Above Mixed | -12.56 | 6.34 | 31.68 | 1.89 | 16.75 |
| Marine Ecology Research Centre | 2/6/2016 | Above Mixed | -9.21 | 4.77 | 16.86 | 1.25 | 13.49 |
|  |  |  |  | Mean | 28.28 |  |  |

Table S2. Clay, silt and sand content, as determined by lazer diffraction (LISST), with their and sediment texture classes at different station across the Salut–Mengkabong lagoon (Sabah, Malaysia). The values are the mean of two replicates at each site and their standard deviations. SD = Standard deviation, mean standard deviation. Mean percentage of silt and clay (SC) and mean percentage of sand (S). STC sediment structure classes

| **Area** | **Station** | **SC** | **S** | **STC** |
| --- | --- | --- | --- | --- |
| Mengkabong | MB | 9.5 ± 6.0 | 94.7 ± 6.0 | Sand |
|  | MC | 8.6 ± 2.8 | 91.4 ± 2.8 | Sand |
|  | MD | 14.4 ± 0.3 | 85.6 ± 14.5 | Sandy–loam |
|  | MF | 21.4 ± 5.8 | 78.7 ± 5.8 | Sandy–clay |
| Salut Lower Branch | SLA | 2.5 ± 0.8 | 97.5 ± 0.8 | Sand |
|  | SLC | 0.9 ± 0.7 | 99.1 ± 0.7 | Sand |
|  | SLE | 9.5 ± 1.0 | 90.5 ± 1.0 | Sand |
|  | SLF | 8.6 ± 11.2 | 91.4 ± 11.2 | Loamy–sand |
| Salut Upper Branch | SUA | 14.5 ± 3.9 | 85.5 ± 3.9 | Sandy–clay–loam |
|  | SUD | 17.9 ± 7.9 | 82.1 ± 7.9 | Sandy–clay–loam |
|  | SUE | 8.9 ± 0.1 | 91.1 ± 0.1 | Silty–loam |
|  | SUF | 8.7 ± 11.1 | 91.3 ± 11.1 | Sand |

Tables S3. a) Seagrass species, aboveground biomass and coverage in Salut–Mengkabong Lagoon. EA: *Enhalus acoroides* and C spp.: *Cymodecea rotundata* and *Cymodecea serrulata.*

| **Sites** | **Area (ha)** | **Station** | **Species** | **Coverage** | **Biomass, t ha^-1^** |
| --- | --- | --- | --- | --- | --- |
| Salut Upper Lagoon | 23.05 | A | EA | ≤ 25% | EA  0.42 ± s.e 0.04 |
|  |  | B | EA | ≤ 25% |  |
|  |  | C | EA | 50 – 75% |  |
|  |  | D | EA | 25 – 50% |  |
|  |  | E | EA | 25 – 50% |  |
|  |  | F | EA | 50 – 75% |  |
| Salut lower lagoon | 7.39 | A | EA, C spp. | Overall ≤50%:  EA ≤ 50%;  C spp. ≤ 25% | C spp.  CR: 0.21± s.e 0.03;  CS: 0.08 ± s.e 0.03 |
|  |  | B | EA, C spp. | Overall ≤75%:  EA ≤ 25%;  C spp. 50 – 75% |  |
|  |  | C | EA, C spp. | Overall ≤75%:  EA ≤ 25%;  C spp. 50 – 75% |  |
|  |  | D | EA, C spp. | Overall ≤50%:  EA ≤ 25%;  C spp. ≥ 50% |  |
|  |  | E | EA | ≥50% |  |
|  |  | F | EA, C spp. | Overall ≤50%:  EA ≥ 50%;  C spp. ≤ 25% |  |
| Mengkabong lower lagoon | 29.04 | A | EA | 25 – 50% |  |
|  |  | B | EA | 50 -75% |  |
|  |  | C | EA | 50 -75% |  |
|  |  | D | EA | 50 -75% |  |
|  |  | E | EA | ≥ 75% |  |
|  |  | F | EA | ≥ 75% |  |

b) Mangrove species, above ground and belowground mean biomass in Salut––Mengkabong Lagoon. RA: *Rhizophora apiculata*; RM: *Rhizophora mucronata*; CD: *Ceriops decandra*, and LR: *Lumnitzera racemosa.* Station 1a: located in upper Salut lagoon (MSL(upper)), which was categorized together with Salut lower lagoon (MSL(lower) because of the similarity of both the sediments, carbon stocks and mangrove species. Station 1b, 2 and 3 were sampled in Salut lower lagoon, near to the lagoon’s entrance (Figure 1).

| Site | Area, ha | Station | Species | Aboveground Biomass, t ha^-1^ | Belowground Biomass,  t ha^-1^ |
| --- | --- | --- | --- | --- | --- |
| Salut upper lagoon | 48.43 | MSU 1 | RA | 392.57 ± s.e 59.90 | 119.43 ± s.e 16.58 |
|  |  | MSU 2 | RA | 291.48 ± s.e 109.06 | 85.59 ± s.e 28.60 |
|  |  | MSU 3 | RA | 90.02 ± s.e 15.99 | 30.86 ± s.e 4.86 |
| Salut lower lagoon | 32.52 | MSL (upper)1a | RA, CD, LR | 287.14 ± s.e 73.87 | 84.98 ± s.e 20.23 |
|  |  | MSL (lower)1b | RA, CD | 793.51 ± s.e 469.66 | 194.53 ± s.e 108.12 |
|  |  | MSL (lower) 2 | RA, RM | 273.99 ± s.e 97.05 | 76.74 ± s.e 24.98 |
|  |  | MSL (lower) 3 | RA, RM, CD, LR | 42.53± s.e 11.12 | 14.49 ± s.e 3.52 |
| Mengkabong lagoon | 68.26 | MMU1 | RA | 186.84± s.e 55.62 | 57.96 ± s.e 15.73 |
|  |  | MMU2 | RA | 299.30± s.e 165.76 | 81.20 ± s.e 38.85 |
|  |  | MMU3 | RA | 259.92± s.e 58.30 | 80.35 ± s.e 16.50 |
| Mengkabong-north shore upper lagoon  (area not sampled) | 681.92 | - | RA | **-** |  |
